# Chronic intermittent fasting impairs β-cell maturation and function in adolescent mice

**DOI:** 10.1101/2024.04.18.590140

**Authors:** Leonardo Matta, Peter Weber, Suheda Erener, Alina Walth-Hummel, Daniela Hass, Lea K. Bühler, Katarina Klepac, Julia Szendroedi, Joel Guerra, Maria Rohm, Michael Sterr, Heiko Lickert, Alexander Bartelt, Stephan Herzig

**Affiliations:** Institute for Diabetes and Cancer (IDC), Helmholtz Center Munich, German Research Center for Environmental Health, Neuherberg, Germany; Institute for Cardiovascular Prevention (IPEK), Faculty of Medicine, Ludwig-Maximilians-Universität München, Munich, Germany; Joint Heidelberg-IDC Translational Diabetes Program, Inner Medicine 1; Heidelberg University Hospital; Heidelberg; 69120; Germany.; German Center for Diabetes Research; Neuherberg, 85764; Germany; Institute of Diabetes and Regeneration Research, Helmholtz Munich, Neuherberg, Germany; School of Medicine, Technical University of Munich (TUM), Munich, Germany; German Center for Cardiovascular Research, Partner Site Munich Heart Alliance, Technische Universität München, Munich, Germany; Chair Molecular Metabolic Control; Technical University Munich; Munich; 80333; Germany

**Keywords:** Intermittent fasting, Langerhans’ islets, beta-cells, insulin, glucose metabolism

## Abstract

Intermittent fasting (IF) is a nutritional lifestyle intervention with broad metabolic benefits, but whether the impact of IF depends on the individual’s age is unclear. Here, we investigated the effects of IF on systemic metabolism and pancreatic islet function in old, middle-aged, and young mice. Short-term IF improved glucose homeostasis across all age groups, without altering islet function and morphology. In contrast, while chronic IF was beneficial for adult mice, it resulted in impaired β-cell function in the young. Using scRNAseq, we delineated that the β-cell maturation and function score were reduced in young mice. In human islets, a similar pattern was observed in Type 1 (T1D), but not in Type 2 diabetes (T2D), suggesting that the impact of chronic IF in adolescence is linked to the development of β-cell dysfunction. Our study suggests considering the duration of IF in younger people, as it may enhance rather than reduce diabetes outcomes.

**Graphical abstract:** 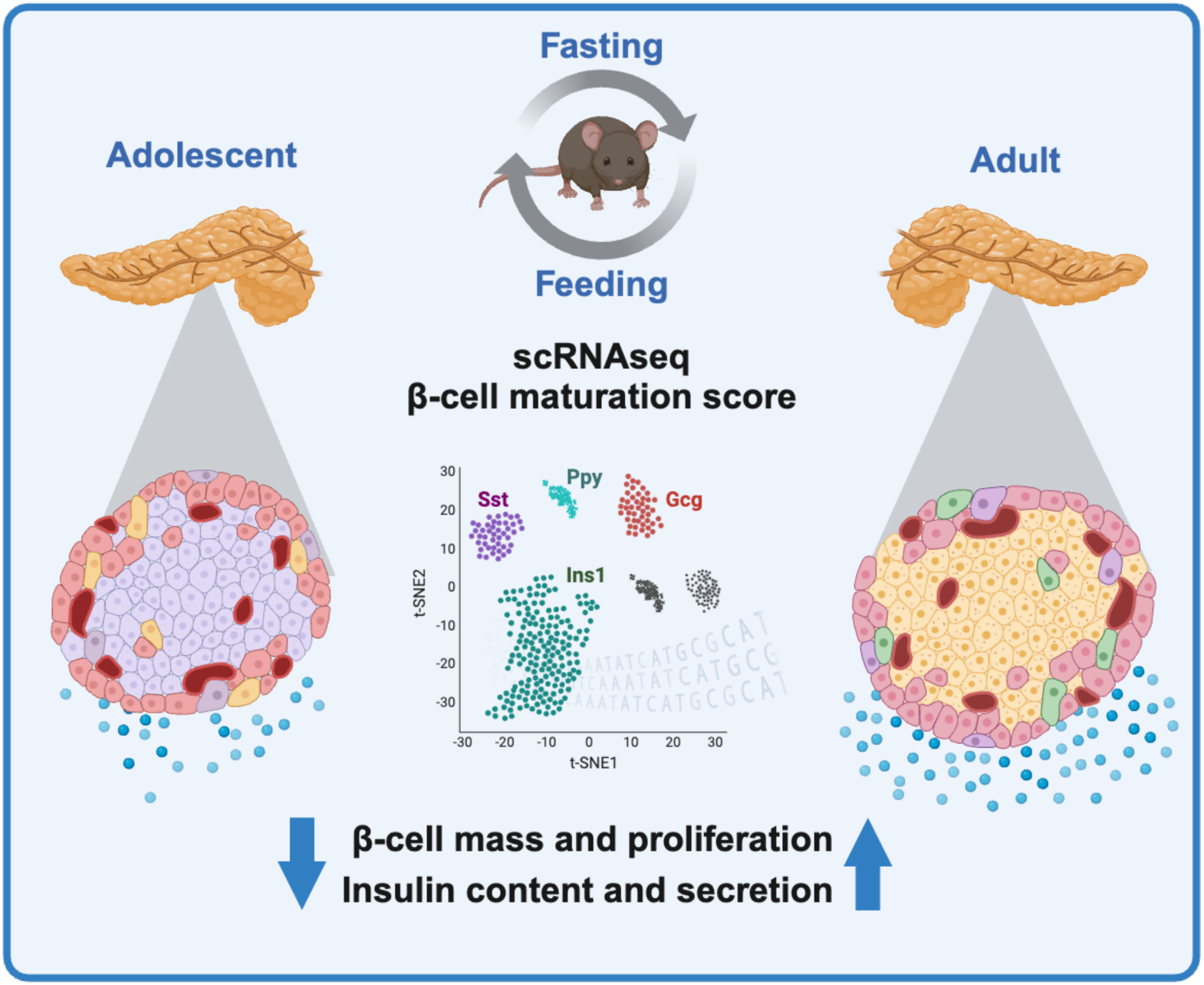

- Long-term IF impairs β-cell function in adolescent mice.
- IF-induced impaired β-cell function is associated with impaired proliferative capacity and lower levels of mature cells.
- IF-induced impaired β-cell function is associated with a transcriptional change highly conserved in type 1-, but not type 2 diabetes.

## Introduction

Modern nutritional habits are characterized by overconsumption of energy-dense and palatable foods, typically rich in lipids and carbohydrates. A positive energy balance, when energy intake exceeds energy expenditure, is associated with the development of obesity and related comorbidities such as diabetes, cardiovascular disease, and cancer^1^. Several nutritional strategies have been explored to prevent weight gain, induce weight loss, and counteract obesity-related diseases to improve metabolic health. Intermittent fasting (IF), a dietary strategy that alternates between periods of fasting and eating, has gained popularity over the past decade. Most human studies suggest that a fasting period of at least ten hours, followed by an eating window for the remainder of the day, is necessary for any metabolic benefits^2^. In mice, alternating between one fasting day and two feeding days leads to improvements in glucose homeostasis, which are largely independent of changes in body weight^3^. Most IF strategies have been proven to exert beneficial effects on metabolic parameters in humans and experimental mouse models, including body weight, insulin sensitivity, plasma cholesterol profiles, blood pressure, and lifespan^4–6^.

The systemic beneficial effects of IF stem from molecular effects on the cellular level. During fasting-feeding cycles, nutrient flux between tissues and cells is stimulated, triggering diverse adaptive responses, including the clearance of potentially detrimental cellular waste. During feeding periods, elevated glucose and insulin levels activate pathways associated with protein synthesis and cell growth, driving cell proliferation and organelle synthesis. Conversely, during fasting, reduced plasma insulin and glucose levels increase cellular energetic stress^7^, which is associated with enhanced cellular turnover and autophagy^8–10^. IF, by reducing cellular glucose flux, decreases adenosine triphosphate (ATP) intracellular levels and increases adenosine monophosphate (AMP). The increase in AMP activates Adenosine monophosphate kinase (AMPK), a sensor of lower energy levels, that also activates PPARG Coactivator 1-α (PGC1-α), stimulating mitochondrial biogenesis and function^11,12^. Enhanced redox homeostasis helps to neutralize ROS, which prevents oxidative stress, cell dysfunction and, chronically, cell death^13^.

These beneficial responses have been implicated in the management of diabetes. Type 2 diabetes (T2D), formerly also known as adult-onset diabetes, is characterized by insulin resistance in the periphery, impaired β-cell function, and hyperglycemia. T2D is associated with β-cell mitochondrial dysfunction, oxidative stress, and inflammation^14,15^. Also, this is associated with blunted growth and proliferative capacity of β-cell, and ultimately results in cell death and loss of β-cell mass^13^. Likewise, Type 1 diabetes (T1D), also referred to as juvenile diabetes, is defined as an autoimmune response that leads to loss of β-cell function and mass^18^. A dysfunctional stress response induced by autoimmune attack and/or viral infection contributes to cell death. In the endoplasmic reticulum (ER), for example, viral infections or inflammatory cytokines induce the accumulation of misfolded proteins and cause ER stress^16^. The pro-inflammatory response is also correlated with mitochondrial impairment and oxidative stress^17^. However, due to the inability of β-cells to properly regenerate, T1D patients display atrophy of β-cells, lower or no levels of intracellular insulin content, and small, highly inflamed islets^18^.

In the context of T2D, IF is associated with increased expression of insulin synthesis and secretion genes, increased insulin content, and improved β-cell function^19^. Besides that, it has been shown that IF protects β-cells by improving antioxidant capacity and decreasing reactive oxygen species formation^19^. In contrast, ^201618^the effects of IF on T1D are largely unclear. Considering that IF promotes tissue regeneration, it is possible that IF influences β-cell dysfunction and death. It has been speculated that IF could provide auxiliary support for individuals with T1D by reducing the need for exogenous insulin. A review of prior studies suggests that fasting in T1D has potential beneficial outcomes when adjustments to insulin are properly managed to support glycemic control^20^. In summary, IF has been proven effective in clinical and pre-clinical settings to counteract metabolic dysfunction observed in T2D and islet dysfunction. However, data on very young and very old subjects are scarce. Based on that, we hypothesize that IF could impact whole-body glucose metabolism, as well as islet function and capacity in an age-dependent manner. To address these hypotheses we employed a combination of immunofluorescence, isolated islets assay, and single-cell RNA sequencing approaches to analyze the effect of IF on glucose homeostasis in mice at various life stages (old, middle-aged, and young).

## Results

### Beneficial metabolic effects of IF in an age-independent manner

IF is a lifestyle intervention that follows various modalities from days to weeks or months. To evaluate the short-term versus long-term effects on metabolic homeostasis and β-cell function, we evaluated the effects of 5 weeks (short-term, ST) and 10 weeks (long-term, LT) IF on body weight and food intake of 2 months old (young), 8 months old (middle-aged), and 18 months old (old) wild-type mice (**Figure 1a**), compared to age-matched control groups fed *ad libitum* (AL). We observed that during the ST-IF intervention, old- and middle-aged mice did not display changes in body weight (**Figure S1a-b and S1f-g**). In the young group on ST-IF, we observed lower body weights, which was linked to lower food intake compared to the young AL control group (**Figure S1k-l**). Furthermore, independent of their age, all groups exposed to the LT-IF intervention displayed lower body weights compared to their respective age control AL groups (**Figure 1b,c**) while food intake remained unchanged (**Figure S1b, g, l**). It is well established that IF has beneficial effects on insulin sensitivity and glucose homeostasis, even independently of weight loss^4^. To this end, we performed dynamic glucose tolerance tests (GTTs) and insulin tolerance tests (ITTs). At the end of the ST-IF intervention, compared to the AL controls, the IF groups exhibited age-independent improvements in both GTT and ITT (**Figure S1c,d, h-j, I)**. Taken together, these results indicate that ST-IF improved glucose homeostasis and insulin sensitivity, independently of age. As the ST-IF intervention improved glucose and insulin tolerance, we monitored if these IF-induced adaptations were sustained in the long term. In old mice subjected to LT-IF, GTT was improved compared to the control AL group (**Figure 1d**). In contrast, the ITT of the LT-IF mice did not differ from the AL group (**Figure 1e**). In middle-aged mice, in comparison to the AL group, LT-IF improved both GTT and ITT (**Figure 1f,g**). To our surprise, the GTT and ITT of young mice were indifferent between the LT-IF and AL groups (**Figure 1h,i**).

**Figure 1.**
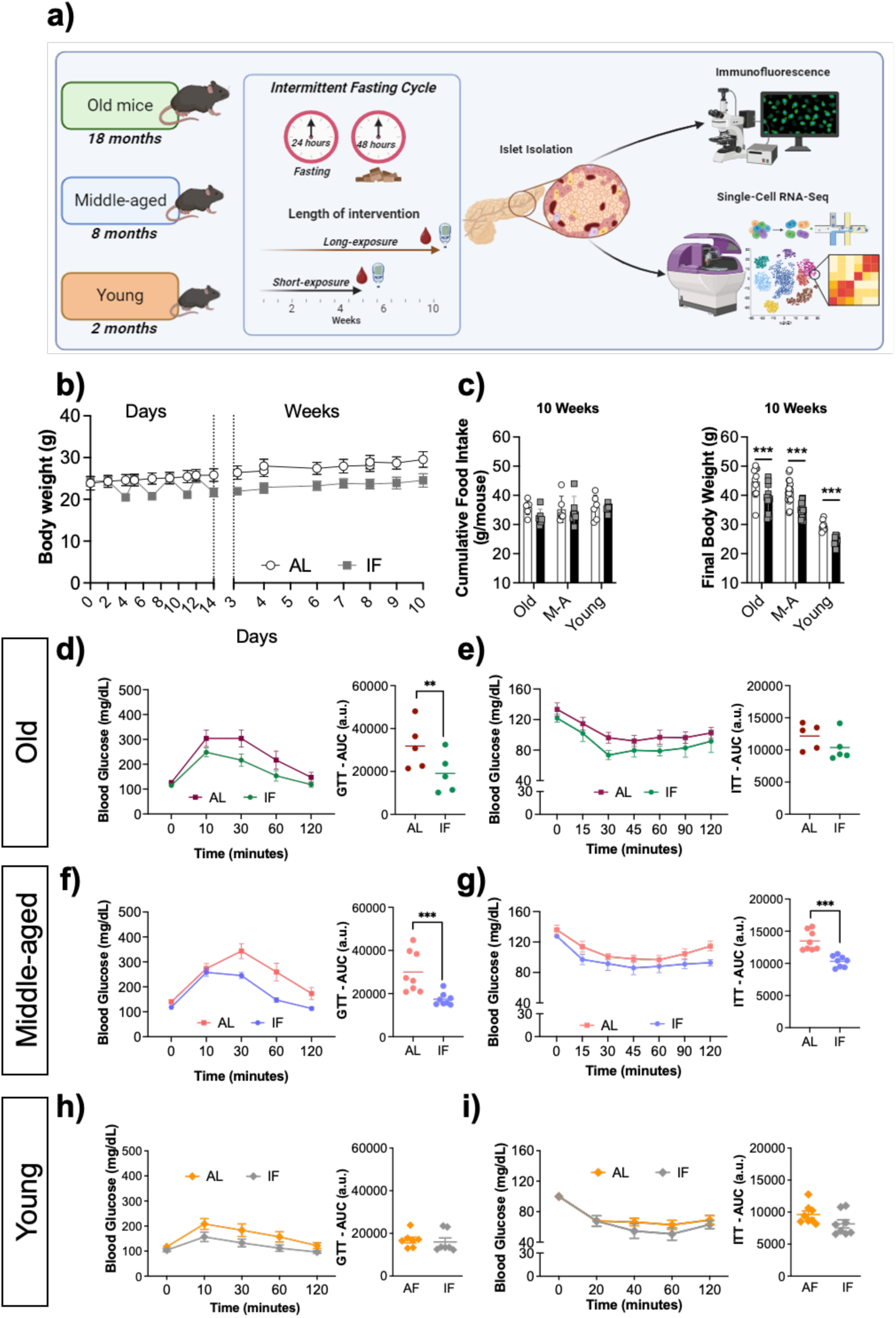
Long-term IF improves glucose homeostasis in middle-aged and old mice, but not in young. Study design from mice *in vivo* and *ex vivo* procedures (a); Daily young body weight during dietary intervention, expressed in days and weeks (b); Cumulative food intake and final body weight 10 weeks of IF intervention (c); Glucose (2.5 mg/kg) and insulin (0.8 U/kg) tolerance tests executed after 10 weeks of IF in old mice (d and e, respectively), middle-aged (f and g, respectively), and young mice (h and i, respectively); Pancreas weight (e, j, and o). AL: Ad libitum; IF: Intermittent fasting; GTT: Glucose tolerance test; ITT: Insulin tolerance test and AUC: Area under curve. The data were expressed as the mean ± standard error of the mean (Old: n = 5/group; middle-aged: n = 8/group; young mice: n= 9/group). Statistical differences, **p<0.01, ***p <0.001.

### Long-term IF impairs β-cells function in younger mice

As one benefit of IF is linked to the endocrine pancreas, we hypothesized that the absence of long-term IF benefits in young mice was potentially linked to β-cells. While, we did not find any difference in pancreas weights between the AL and LT-IF groups, independently of age (**Figure S1e,j,o**) we investigated islet and β-cell function further. In isolated primary islets, we performed glucose-stimulated insulin secretion (GSIS) tests to directly assess β-cell function. At low glucose concentrations, islets from the LT-IF and AL groups secreted comparable insulin levels. However, islets isolated from old LT-IF mice secreted more insulin in response to high glucose than islets isolated from the AL group (**Figure 2a**), while no significant changes were observed in islets isolated from the middle-aged mice (**Figure 2b**). Contrary to this, islets isolated from young mice after LT-IF showed lower GSIS compared to the control AL group (**Figure 2c**). Maximal secretion capacity, as assessed by KCl-induced insulin secretion, was indifferent in any group, although it followed the pattern observed for GSIS (i.e., a trend towards higher secretion in old and lower secretion in young IF mice) (**Figure 2a-c**).

**Figure 2.**
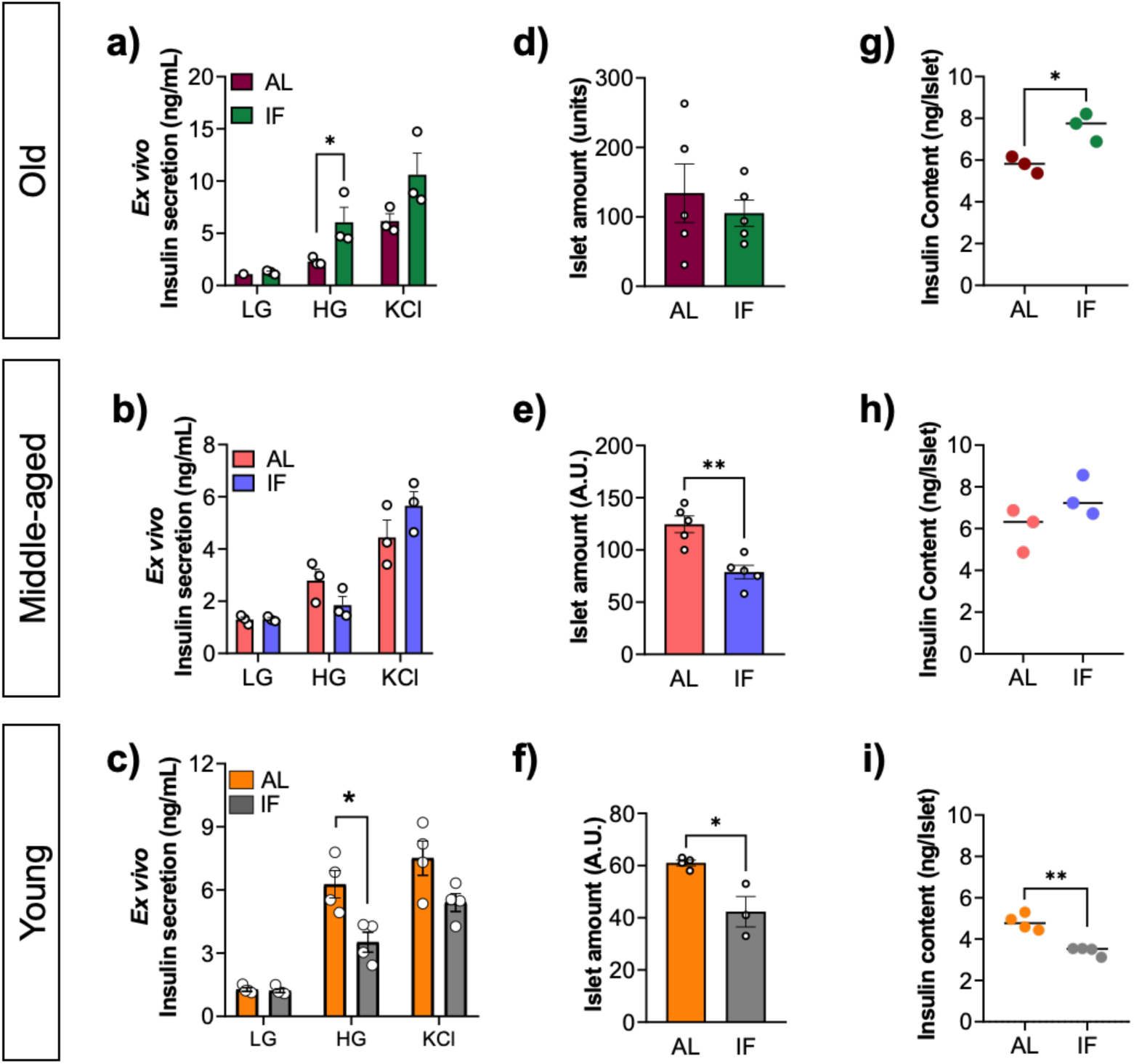
Long-term IF increases insulin content and secretion in old-, but decreases in young mice. *Ex vivo* insulin secretion, islet amount, and insulin content after 10 weeks of IF in old mice (a, b, and c, respectively), middle-aged (d, e, and f, respectively), and young mice (g, h, and i, respectively). AL: Ad libitum; IF: Intermittent fasting. The data were expressed as the mean ± standard error of the mean (Old: n = 3/group; middle-aged: n = 3/group; young mice: n= 4/group) for islet amount measurement (Old: n = 4/group; middle-aged: n = 5 - AL group and 4 IF group; young mice: n= 3/group). Statistical differences, *p<0.05, **p <0.01.

To explore the mechanism of these age-dependent effects of LT-IF on insulin secretion, we quantified islet amount and insulin content. Interestingly, the islet quantity was unchanged in the old mice (**Figure 2d**) but lower in the middle-aged (**Figure 2e**) and young LT-IF groups compared to their AL controls (**Figure 2f**). The insulin content of isolated islets was higher in the old (**Figure 2g**), did not change in the middle-aged (**Figure 2h**), and was lower in the young LT-IF group (**Figure 2i**) compared to their AL control groups. Altogether, our data indicated that LT-IF improved *ex vivo* islet function in old mice and impaired islet function in adolescent mice.

### Impaired islet function in young mice is linked to lower α/β-cell mass and proliferation

Based on the impaired islet function after LT-IF in young mice, we next investigated whether IF affected α- and β-cell mass and proliferation. Mice were injected with BrdU^+^ for 5 days after 10 weeks of IF. By immunofluorescence, we observed that although there was no difference in islet area (sum of β-, α-cell populations and nuclei) (**Figure 3a-b**), islets in young mice subjected to long-term IF had lower α- (**Figure 3c**) and β-cell masses (**Figure 3d**), and α/β- cell ratios compared to the AL group (**Figure 3e**). Additionally, we observed that LT-IF was associated with less BrdU^+^ labeling in insulin and glucagon-positive cells, pointing to reduced proliferation of both α- and β-cells (**Figure 3f-g**). In old mice, quantification for insulin and glucagon-positive cells indicated that compared to the AL control group, islets from LT-IF had lower α-cell mass (**Figure S2a-c**). However, islet area, β-cell, BrdU-positive insulin, and glucagon-positive cells were indifferent between the AL and LT-IF groups in old mice (**Figure S2b-g**). In middle-aged mice, no significant changes in islet area, α- and β-cell mass, α/β-cell mass ratio, and α- and β-cell proliferation staining were observed between the groups (**Figure S2h-n**). Taken together, our data suggest that in young mice, LT-IF impaired α- and β-cell proliferation, contributing to reduced islet function and insulin secretion. However, this was not seen in old- and middle-aged mice.

**Figure 3.**
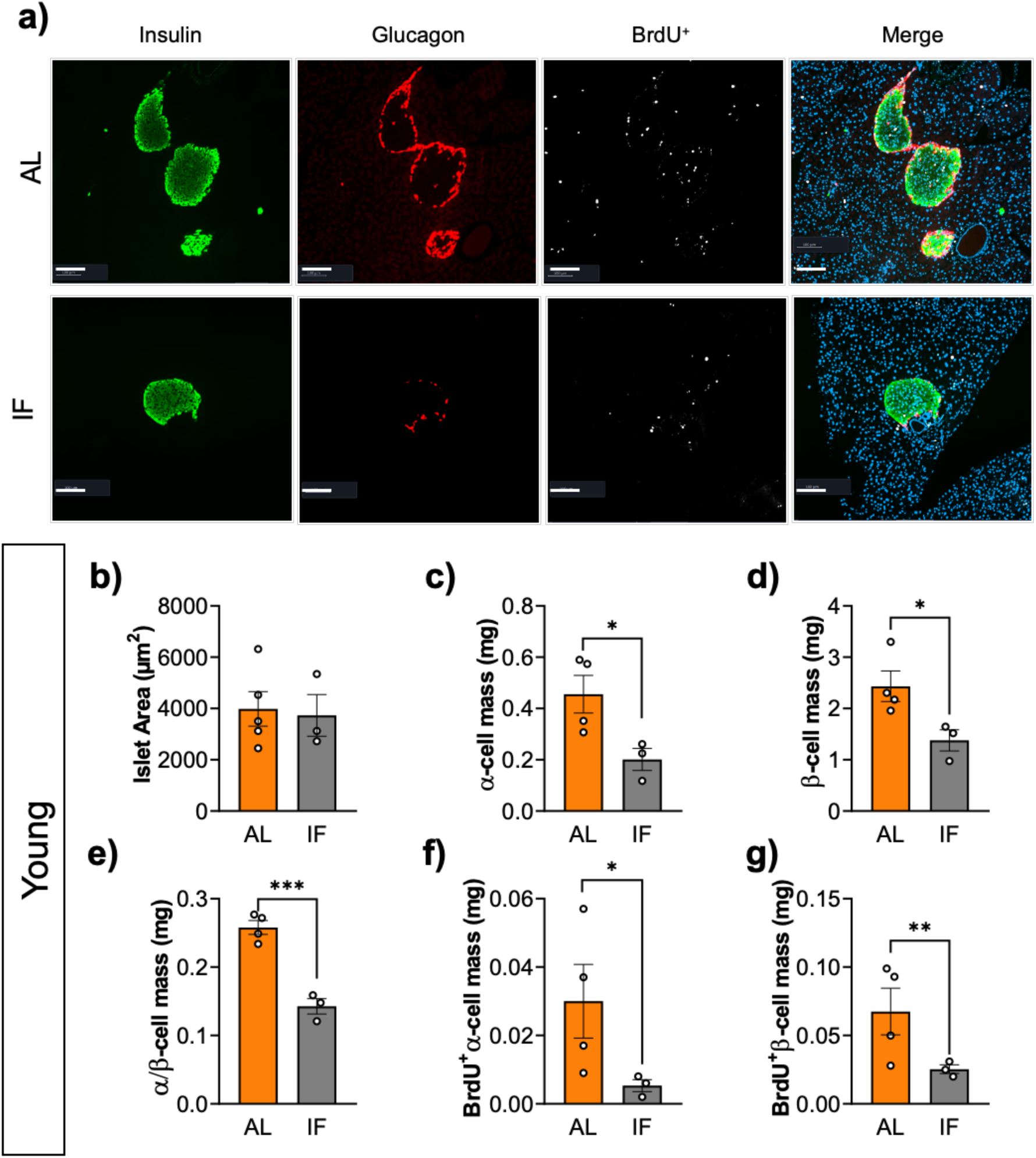
Lower insulin secretion and content in young IF mice are associated with reduced islet numbers and impaired α- and β-cell proliferative capacity. Immunofluorescence from isolated islets from young mice after long-term IF (a); Islet area μm2 (b); α-cell mass (c); β-cell mass (d); α- and β-cell ratio (e); α- and β-cell mass - positively stained for BrdU – proliferation (f and g, respectively). Insulin positive cells: Green; Glucagon positive cells: Red; BrdU positive cells: White; and nuclear staining (DAPI): Blue – Scale bar 100μm. AL: Ad libitum; IF: Intermittent fasting. The data were expressed as the mean ± standard error of the mean (n=4 in the AL group and 3 in the IF group). Statistical differences, *p<0.05, **p <0.01, ***p<0.001.

### Islet scRNA-seq reveals altered β-cell maturation and function in young LT-IF mice

In old mice, LT-IF was associated with enhanced islet function but was impaired in young mice compared to their AL controls. To unravel the underlying differences in function and cell populations, we performed scRNA-seq of isolated islets from young, middle-aged, and old mice after LT-IF **(Figure 4a)**. From the expression matrix with cells as columns and genes as rows, we assigned cell clusters to the main endocrine cell types based on the established marker genes *Gcg* (encodes for glucagon found in α-cells), *Ins1* (encodes for insulin found in β-cells), *Ppy* (encodes for pancreatic polypeptide found in PP cells), and *Sst* (encodes for somatostatin in δ-cells) (**Figure S3b**). After duplex removal of the main endocrine cell types, we distinguished distinct *Gcg, Ins1, Ppy,* and *Sst*-positive cell populations (**Figure 4b,c**). In addition, we found the expected non-endocrine cell types, confirming the validity of our scRNA-seq approach (**Figure S3c**). Further unsupervised cell clustering from the whole data set revealed 22 sub-cell populations (**Figure S3a**). We focused our bioinformatic analyses on β-cells to determine age and IF-dependent transcriptomic responses (**Figure 4d**, **Figure S3d,e; Table S1**). We performed a quantitative analysis of islet-specific biological programs described in the literature^21^, which were augmented with custom gene sets (**Table S1**). Module scores based on those programs resemble features that can be interpreted functionally and are tailored to the specific biological context in question. We found that the most significant difference between AL and LT-IF groups was the “beta cell maturation” genes set (**Figure 4e, Figure S3g, Table S2)**. Whereas in old and middle-aged mice β-cell maturation scores were higher after LT-IF, they were lower in young mice compared to their AL controls (**Figure 4e)**. Also, when we stratified this analysis by heterogenous β-cell clusters we found the same effects in all clusters (**Figure S3g**). We also investigated the overrepresentation of KEGG pathways in the differentially regulated genes of β-cells (**Table S3**). We identified a cluster of related KEGG pathways that included pathways for T2D and insulin function (**Figure 4f, Figure S3f)**. We used the overlap of genes annotated to the “beta cell maturation” program with those contributing to any pathway overrepresentation of the found clusters to identify possible candidate genes, as highlighted in volcano plots (**Figure 4d, Figure S3d,e**) and visualized as a heatmap (**Figure S3h**). Most of these candidate genes such as *Mafa, Nkx-6-1, Slc2a2,* and *Ins1* were lower only in young mice subjected to LT-IF compared to AL but not in middle-aged or old mice (**Figure S3h**). Next, we further applied external validation and translation of the candidate genes into human data by interrogating published datasets from human islets of T2D and T1D donors. We observed that our candidate genes, which were specifically lower in the young-IF group, showed a similar pattern of downregulation in T1D, but not in T2D (**Figure 4g,h)**. In conclusion, our scRNA-seq analysis revealed a specific β-cell gene signature in young mice subjected to LT-IF, which was linked to T1D in humans.

**Figure 4.**
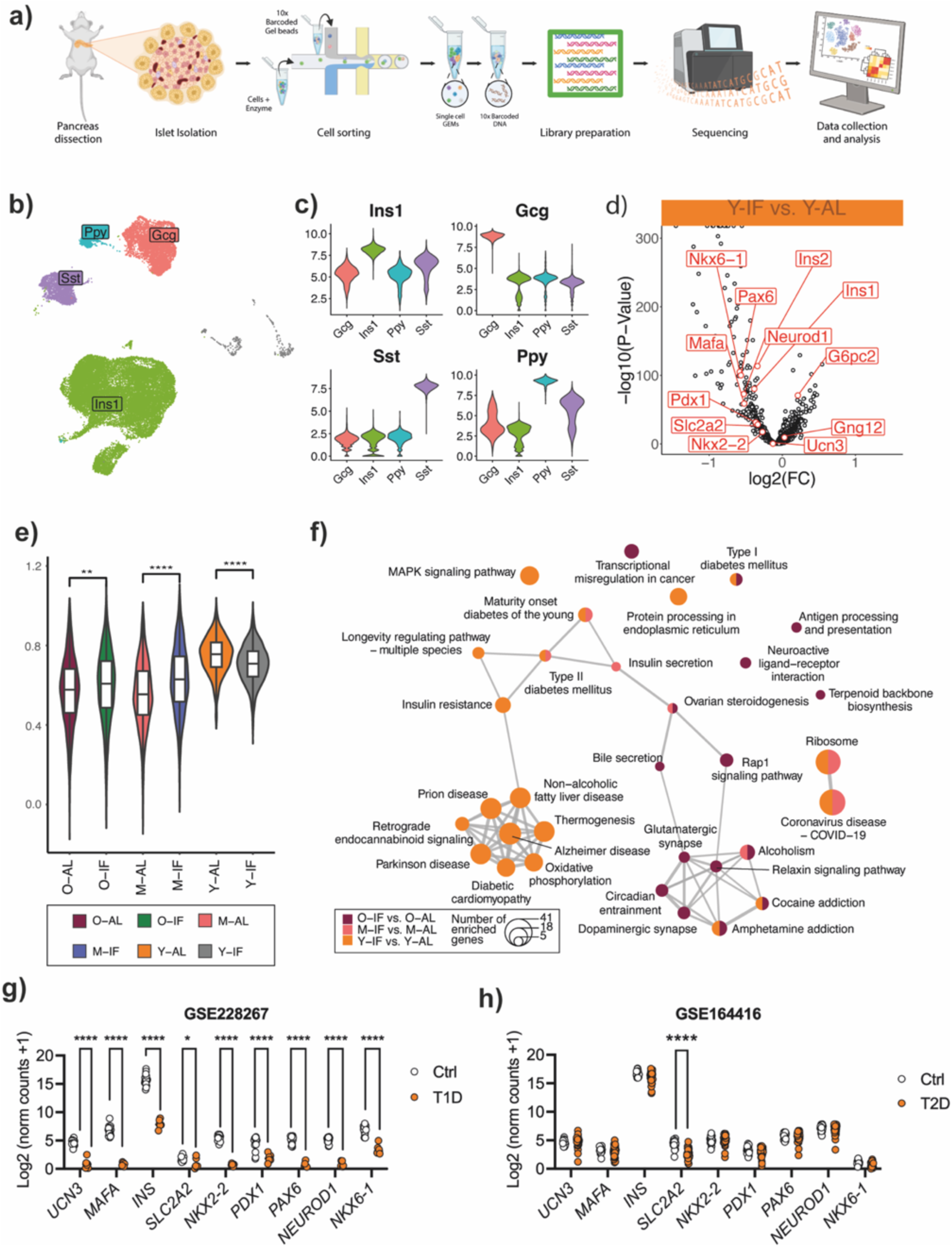
scRNA-seq reveals impairs β-cell maturation and development in young mice. scRNA-seq processing in isolated islet (GEM = Gel bead in emulsion) (a). UMAP of whole data set after duplex removal with color-coded endocrine cell types; labels show main marker genes (b) and violin plots showing expression of marker genes in the identified cell types (c). Differential gene expression is visualized as a volcano plot with biological effect size (log2(FC) on the x-axis and statistical significance (-log10(P-value) on the y-axis (d). Comparison of module scores for beta cell maturation markers between IF and AL in all age groups (e) Statistical significance * padj<0.05; ** padj<0.01; *** padj<0.001; **** padj< 0.0001). Overrepresentation of KEGG pathways from differentially expressed gene colors indicates contrasts in which significant results were found: Intermittent fasting vs ad libitum group in old (O-IF vs. O-AL), middle-aged (M-IF vs. M-AL), and young mice (Y-IF vs. Y-AL); connected clusters are based on semantic similarity and size of nodes indicate several genes (f). *In silico* validation of differential gene expression in the isolated β-cells from human donors with type I diabetes (g) or type II diabetes (h). Statistical differences,*p<0.05, ****p<0.0001.

## Discussion

Despite the existing evidence of the beneficial effects of IF on metabolic health, if these apply to all life stages remain largely unexplored. Most IF studies were understandably conducted in adult humans or middle-aged/elderly mice, as these are at higher disease risk compared to the young. Still, there is limited knowledge about the impact of IF on younger populations^22^. In this study, we investigated the impact of age on IF-induced improvements in glucose homeostasis. We found that while ST-IF had metabolic benefits across all tested age groups, LT-IF negatively impacted pancreatic islet function only in young mice. Using scRNA-seq of isolated islets, we discovered a transcriptional profile linked to dysfunctional β-cell maturation specifically in the young. Further bioinformatic analysis revealed that this signature was linked to a transcriptional signature of β-cell maturation and function-related genes observed in isolated islets from humans with T1D.

Overall, our data align well with previous findings, demonstrating that a short IF regimen enhances insulin sensitivity^4,5,8^. The finding that longer IF periods may also have deleterious consequences is a novel finding with relevant implications, especially for using IF in adolescents and people at high T1D risk. Previous studies have reported that a fine balance in nutrient flux is necessary for tissue maturation and function, and fasting interventions in immature mice may trigger metabolic impairment. In agreement with our findings, Yin, Wenzhen, et al. (2023) demonstrated that 12 weeks of IF intervention, in pregnant mice, predisposes their offspring to liver steatosis and adiposity, suggesting that IF should be avoided during pregnancy due to its potential adverse effects on fetal development and offspring metabolic homeostasis^23^. Bayliak, M. et al. (2021) found that while IF had beneficial effects on redox homeostasis and energy metabolism in the older groups, it appeared to decrease antioxidant capacity in the cortex of young mice^24^. These findings align with the results from Jianghua, et al. (2011), which showed that IF affects brain energy metabolism by reducing brain insulin signaling and, consequently, peripheral insulin production and secretion^25^. Taken together, also previous studies indicate a potential impairment induced by long-term IF interventions. In light of our study, this supports the notion that during the period of development and maturation, IF might impair proper nutrient flux and hormonal balance required for proper cell differentiation and organ development.

The young mice displayed reduced α- and β-cell mass after LT-IF, which was not observed in the middle-aged or old mice. Reduction in cellular nutrient flux may lead to an atrophic cellular remodeling. Atrophic responses are generally associated with secretory malfunction^26,27^, an early indication of β-cell failure^28,29^, including altered pro-insulin processing, elevated pro-insulin levels, a reduction in secretory granules, and β-cell deterioration^27,28^. These pathological responses can be triggered by genetic predisposition (lowered secretory capacity) and environmental stressors. Imbalances in calcium-, redox- or mitochondrial homeostasis lead to a pro-oxidative and -inflammatory environment and, in later stages to β-cell apoptosis^29^. Furthermore, suboptimal nutrient flux may contribute to impaired cellular maintenance and development^30^. Although the direct effects of these factors on the secretory program of immature β-cells remain unclear, it might very well be related to the deleterious effect of IF observed in our study^32^.

Understanding environmental factors influencing β-cell development, function, and failure is an important step toward enhancing the prediction, diagnosis, and treatment of diabetes. Our scRNA-seq analyses revealed a distinct molecular signature following LT-IF in younger mice. The key findings include the higher proportion of genes indicating immature β-cells and downregulation of maturation- and β-cell function-related genes, particularly *Mafa, Nkx6-1, Slc2a2,* and *Ins1*. The products of these genes are important for glucose transport as well as insulin synthesis and secretion, potentially hinting at the impact and mechanisms of IF on β-cells. Our data set adds a new perspective on the transcriptional modulation associated with impaired β-cells architecture and function. The data indicate a potential vulnerability of developing β-cell when subjected to extended fasting periods, which raises caution for extended fasting in individuals at risk of developing diabetes. In T1D, where β-cell preservation is paramount, IF may exacerbate the underlying β-cell fragility. This is particularly relevant given the increasing popularity of IF as a lifestyle choice among diverse age groups. An interesting finding is that we also performed an in-silico analysis of public data and observed the same transcriptional patterns in humans with T1D, which was not observed in T2D. However, it remains to be seen how our findings translate into settings where high-fat diet-induced obesity is the main driver of islet dysfunction or in models of T1D such as NOD mice.

In conclusion, our study underscores the importance of tailored approaches in dietary practices and highlights the need for comprehensive longitudinal studies to elucidate the long-term effects of IF, particularly in human populations and individuals with a predisposition to diabetes. While IF exhibits promising metabolic benefits in the short term, the potential age-dependent adverse effects observed with prolonged fasting require more attention and caution.

### Limitations of the study

This study provides valuable insights into the age-dependent effects of IF on glucose metabolism and pancreatic islet function in mice. While our findings are primarily derived from a mouse model, which, despite its usefulness in preclinical research, cannot completely emulate human metabolic processes. The age classifications used correspond to mouse physiology and may not translate directly to human life stages, where developmental and metabolic processes differ significantly. Lastly, the focus of our study on pancreatic function and glucose homeostasis means that other physiological or behavioral effects of IF, such as those related to cognitive function or physical activity, were not assessed. Future studies should address these limitations by incorporating a broader range of IF practices, extending the duration of the study, and including a more diverse set of physiological, once just male mice were used, as well as behavioral endpoints.

## Acknowledgments

We thank all lab members for the discussions and the enjoyable atmosphere. S.H. was supported by the Helmholtz Future Topic AMPro. A.B. was supported by DFG (SFB1123-B10 and SPP2306 BA4925/2-1), the Deutsches Zentrum für Herz-Kreislauf-Forschung (DZHK), and the European Research Training Group (ERC) Starting Grant PROTEOFIT. M.R. is funded by the European Research Council (ERC) under the European Union’s Horizon 2020 research and innovation program (# 949017) and the German Diabetes Center (DZD) Next grant. We acknowledge BioRender.com for help with the figures. We apologize to colleagues whose work we could not cite due to space limitations.

## Author contributions

**L.M.**: Conceptualization; data curation; formal analysis; visualization; writing – original draft, review, and editing. **P.W.**: Conceptualization; data curation; formal analysis; visualization; methodology; writing – scRNA-seq methodology, review, and editing. **S.E.**: Investigation, project administration, conceptualization; data curation; formal analysis; visualization; methodology; writing – review. **A.W-H.**: Investigation; visualization; methodology; writing – review and editing. **D.H.**: Investigation; visualization; methodology; writing – review and editing. **L.K.B.:** Investigation; visualization; methodology; writing – review. **K.K.**: Investigation; visualization; methodology; writing – review. **J.S.:** Investigation; visualization; methodology; writing – review. **J.G.:** Data curation; formal analysis; visualization; methodology; writing – review. **M.R.**: Investigation, project administration, conceptualization; data curation; formal analysis; visualization; methodology; writing – editing, and review. **M.S.**: Data curation; formal analysis; visualization; methodology; writing – review. **H.L.**: Data curation; formal analysis; visualization; methodology; writing – review. **A.B.**: Conceptualization; supervision; writing – original draft; project administration; writing – review and editing. **S.H.**: Conceptualization; supervision; funding acquisition; writing – original draft; project administration; writing – review and editing.

## Methods

### Key Resources Table

| REAGENT or RESOURCE | SOURCE | IDENTIFIER |
| --- | --- | --- |
| **Antibodies** |  |  |
| Insulin (C27C9) Rabbit mAb | Cell Signaling | Cat# 3014. RRID: AB_2126503 |
| Anti-Glucagon Mouse mAB | Sigma-Aldrich | Cat# G2654. RRID: AB_259852 |
| Donkey Anti-rabbit Antibody, Alexa Fluor 594 Conjugated | Molecular Probes | Cat# A-21207. RRID: AB_141637 |
| Donkey Anti-mouse Antibody, Alexa Fluor 488 Conjugated | Molecular Probes | Cat# A-21202. RRID: AB_141607 |
| 4', 6-diamino-2-phenylindole - DAPI | Vector Laboratories | Cat#H1200. RRID: N/A |
| Mouse Anti-BrdU Monoclonal Antibody | Abcam | Cat# ab8152. RRID: AB_308713 |
| Goat polyclonal Secondary Antibody to Mouse IgG2b - (FITC) | Abcam | Cat# ab98702. RRID: AB_10696531 |
| Goat anti-Mouse IgG H&L (Cy3) secondary antibody | Abcam | Cat# ab97035. RRID: AB_10680176 |
| **Bacterial and virus strains** |  |  |
| NOT APPLIED |  |  |
| Biological samples |  |  |
| NOT APPLIED |  |  |
| **Chemicals, peptides, and recombinant proteins** |  |  |
| Roswell Park Memorial Institute (RPMI) Medium | Thermo Fisher | Cat# 1640 / RRID: N.A |
| Krebs-Ringer Solution, HEPES-buffered | Thermo Fisher | Cat# J67795.K2 RRID: N/A |
| **Critical commercial assays** |  |  |
| Ultra Sensitive Mouse Insulin ELISA Kit | Crystal Chem | Cat#90080 . RRID: N/A |
| Histofix ® 4% | Roth | Cat# P087.3 . RRID: N/A |
| **Deposited data** |  |  |
| Raw and analyzed data – Figure 4g | This paper | GSE228267 |
| Raw and analyzed data – Figure 4h | This paper | GSE164416 |
| **Experimental models: Cell lines** |  |  |
| NOT APPLIED |  |  |
| **Experimental models: Organisms/strains** |  |  |
| Male mice – Strain: C57BL/6N | Janvier Laboratory | RRID:MGI:2159965 |
| **Oligonucleotides** |  |  |
| NOT APPLIED |  |  |
| **Recombinant DNA** |  |  |
| NOT APPLIED |  |  |
| **Software and algorithms** |  |  |
| Prism (GraphPad) Version 9 |  | https://www.graphpad.com/ |
| ImageJ |  | https://imagej.nih.gov/ij/ |
| The R Project for Statistical Computing v 4 | CRAN | https://www.r-project.org/ |
| CellRanger v2 & v3 | 10x Genomics | https://www.10xgenomics.com/support/software/cell-ranger/latest |
| Seurat v 3 | CRAN | https://satijalab.org/seurat/articles/install_v5 |
| DESeq2 | Bioconductor | DOI: 10.18129/B9.bioc.DESeq2 |
| ggplot2 | CRAN | https://ggplot2.tidyverse.org/ |
| ComplexHeatmap | Bioconductor | DOI: 10.18129/B9.bioc.ComplexHeatmap |
| Other |  |  |
| ACCU-CHEK Performa | Roche | Cat#: 9055599 . RRID: N/A |

### Experimental model

The study was approved by the government of Upper Bavaria, according to the protocol number: ROB-55.2-2532. Vet_02-19-256, and performed by the Laboratory Animal Resources guidelines. Five-week-old male C57BL/6N mice were purchased from Janvier Laboratory. The colony was housed in groups of four, in a specific pathogen-free (SPF) facility in individually ventilated cages and a controlled environment (21-22°C, 30%-60% humidity) with free access to water and a standard rodent chow diet (18.9% protein, 5.5% fat, and 61.3% carbohydrate). All mice were subdivided into three groups, to evaluate the effect of IF at different stages of life; the old group (eighteen months old); the middle-aged (eight months old), and the young (two months old).

### Intermittent fasting

Old, middle-aged, and young mice were randomly grouped into ad libitum (AL) and intermittent fasting (IF) groups. In the IF group, the food was removed from 10 a.m. to the following day (24 hours later), we employed the intermittent fasting protocol 1:2, which consists of 24 hours of food starvation for 48 hours of free access to a chow diet, an adaptation from protocols previously described and studied ^31–33^. Mice in the AL group were handled similarly but with unrestricted chow diet access. For all groups, water was offered without restriction. Food intake was measured every three days, in the IF group we calculated the difference between the amount of food offered at the beginning of the food offer phase with the rest at the end of the phase. All groups, independently of age, were exposed to a period of 4-5 weeks (11 IF cycles) or 9-10 weeks (22 IF cycles) of the IF regimen and euthanized by cervical dislocation forty-eight hours after the last IF cycle.

### Glucose tolerance tests (GTT) and insulin tolerance tests (ITT)

Mice were fasted for 6 for GTT, and a hypertonic solution of glucose was administered intraperitoneally (2.5 g/kg). For ITT, we fasted the mice for 5 hours (morning fasting) and insulin (0.8 units/kg) was injected intraperitoneally. Both tests measured blood glucose levels at 0, 15, 30, 60, 90, and 120 minutes after each injection. Blood glucose was measured using an Accu-Chek Performa glucose monitor (Roche Diagnostics, Castle Hill, Australia). Plasma insulin was measured using ELISA (Crystal Chem, Downers Grove, IL)^34^.

### Islet Isolation

Mice were sacrificed by cervical dislocation. The peritoneal cavity was exposed for clamping of the pancreatic main duct to the intestine and cannulation in situ via the common bile duct. Pancreata were inflated with a Collagenase P solution (2 U/ml in Hank’s buffered salt solution (HBSS) supplemented with 1% BSA) and dissected, followed by an incubation in a 37°C water bath for 14 minutes. Islets were hand-picked four times in HBSS supplemented with 1% BSA, cultivated in Roswell Park Memorial Institute (RPMI) media containing 5.5 mmol/L glucose, 10% fetal bovine serum (FBS), and 1% penicillin and streptozotocin solution, and incubated at 37°C and 5% CO2^35^.

### Insulin secretion and content assay

20 islets, with three replicates per animal, were washed twice in Krebs-Ringer HEPES buffer (containing 118.5 mM NaCl, 25 mmol/L NaHCO3, 1.19 mmol/L KH2PO4, 2.54 mmol/L CaCl2, 1.19 mM MgSO4, 4.74 mmol/L KCl, 10mmol/L HEPES, and 0.1% BSA) supplemented with 1 mmol/L glucose. Islets were then starved in Krebs-Ringer HEPES Buffer containing 1 mmol/L glucose for 1 hour at 37°C. Insulin secretion was stimulated at low glucose levels (LG - 2.8 mmol/L glucose), followed by high-glucose levels (HG - 16.7 mmol/L glucose), and potassium chloride (KCl - 20 mmol/L) for 30 min at 37°C. Islets were harvested in acid ethanol (1.5% HCl in 70% ethanol) to determine insulin content. Insulin levels were determined by ELISA following the manufacturer’s instructions (Crystal Chem, Cat#90080, Downers Grove, IL). Insulin secretion and content were normalized to islet numbers^36^.

### Immunofluorescence

The pancreas was dissected and fixed with Histofix^®^ (paraformaldehyde 4%) for sections. Following, short bouts of microwave exposure were used for antigen retrieval in a solution containing 1 mM EDTA, pH 8.0, for 15 minutes, followed by blocking in 1% BSA (Santa Cruz Biotechnology, sc-2323), 4% normal donkey serum (Abcam, AB166643) for one hour at room temperature. Indirect immunofluorescence was performed using primary antibodies, rabbit anti-insulin (1:100; Cell Signaling Technology, Cat# 3014. RRID: AB_2126503), mouse anti-glucagon (1:500; Sigma-Aldrich, Cat# G2654. RRID: AB_259852); and secondary antibodies, namely, Alexa Fluor 594 donkey anti-rabbit (1:500) and Alexa Fluor 488 donkey anti-mouse (1: 500; Molecular Probes, Cat# A-21207. RRID: AB_141637, and Molecular Probes, Cat# A-21202. RRID: AB_141607, respectively). Nuclei were stained with 4′, 6-diamino-2-phenylindole (DAPI; Vector Laboratories, H1200). Images were acquired using a Zeiss Confocal LSM 700 Laser Scanning confocal microscope using 40X Zeiss Plan-Neofluar 40X/1.3 and 63X/1.4 oil immersion objectives. The β-cell area was assessed by ImageJ^37^.

### BrdU incorporation and detection

The procedure for BrdU labeling was previously described by Téllez, Noèlia, and Eduard Montanya^38^. Briefly, an analog of thymidine, BrdU (100 mg/kg body weight), was injected intraperitoneally twice a day for 5 days. For BrdU detection, the pancreas was collected and fixed in paraformaldehyde 4% before staining. The fixed pancreas slides were incubated with 0.5 M HCl at 37°C for 30 minutes for DNA denaturation. The slides were washed four times with PBS. Mouse anti-BrdU antibody (1:100; Abcam, Cat# ab8152. RRID: AB_308713) was applied at 4°C for 16 hours. A FIT-conjugated goat anti-mouse (1:10, Abcam, Cat# ab98702. RRID: AB_10696531) or Cy3-conjugated sheep anti-mouse IgG (1:200, Abcam, Cat# ab97035. RRID: AB_10680176) was used to reveal the BrdU incorporation.

### Single-cell RNA-Sequencing Sample Preparation and Data Generation

For scRNA-seq, islets viability was measured and posteriorly dissociated using TrypLE Express (Gibco) for 12 to 20 minutes at 37 °C. Cells were then washed with PBS containing 2% FCS, filtered through a 40 μm Flowmi™ Cell Strainer (Bel-Art), and diluted to a final concentration of ∼1000 cells/µl in PBS containing 2% FCS. The cell suspension was immediately used for scRNA-seq library preparation with a target recovery of 10000 cells. Samples from young mice; MUC8408, MUC8409, MUC8410, MUC8411, MUC8412, MUC8413: Libraries were prepared using the Chromium Single Cell 3ʹ Reagent Kits v2 (10x Genomics) according to the manufacturer’s instructions. Libraries were pooled and sequenced on an Illumina HiSeq4000 with a target read depth of 50000 reads/cell. FASTQ files were aligned to the mm10 mouse genome with Ensembl release 94 annotations and pre-processed using the CellRanger software v2.1.1 (10x Genomics) for downstream analyses. Samples from middle-aged and old mice; MUC28169, MUC28170, MUC28171, MUC28172: Libraries were prepared using the Chromium Single Cell 3ʹ Reagent Kits v3.1 (10x Genomics) according to the manufacturer’s instructions. Libraries were pooled and sequenced on an Illumina NovaSeq6000 with a target read depth of 50000 reads/cell. FASTQ files were aligned to the GRCm38 mouse genome with Ensembl release 101 annotations and pre-processed using the CellRanger software v3.1.0 (10x Genomics) for downstream analyses.

### ScRNA-seq data processing, quality control, and analysis

Preprocessed count data were quality controlled with scatter 1.10.1^39^, imported into and analyzed by Seurat (v3)^40–42^. Filtering of cells was based on more than 1500 but less than 8000 expression features in young mice and between 1500 and 5500 in the old group; for both mitochondrial fractions of reads < 10%. For normalization and variance stabilization, we used the modeling framework sctransform^43^, following the Seurat v3 vignette “vignettes/sctransform_vignette.Rmd”. Integration of individual samples has been performed using Pearson residuals as described in Seurat’s SCTtansform workflow. Dimensionality reduction plots are UMAP built with 30 PCs and confound removal was based on mitochondrial percentage after integration. Identification of statistical clusters was performed before duplex removal with Seurat’s internal function “FindClusters” with the parameter resolution = 0.5. It is based on a shared nearest neighbor (SNN) modularity optimization method by Waltman L., and Van Eck N.^44^ Cell identity classification and duplex removal: The canonical marker gene expression for the main endocrine cell types was extracted from the “RNA” slot and thresholded based on individual inspection of the data distributions (“Ppy”>8.5, “Gcg” >8, “Sst”>4, “Ins1”>8.7). Cells with high expression of exclusively one marker were used as pure profile cells in further analysis. Conserved marker genes have been identified for each of the four cell types from the pure profile cells and subsequently, centroids per cell type have been computed comprising all conserved marker genes from any of the pure profile cell types. In addition, artificial mixed-type centroids were computed for each pairwise and triple-cell combination. In the final step, transcriptomic profiles of all cells were individually correlated to all (pure and mixed) centroids and classified according to the highest correlation. Cells with the highest correlations to mixed centroids were finally removed as duplets. Quality control of classification was based on plausibility check in UMAPs, marker expression, and comparison to similar results obtained with the package “Scrublet”, as duplet removal^45^. Differential gene expression analysis was done using DEseq2 statistical models from the RNA assay with the function “FindMarkers” and min.pct=0.3^46^. Since there are no biological replicates we used individual cells for computing inference statistics. Genes with p_val_adj<0.1 & abs(avg_log2FC) >0.2 were used for over-representation-based pathway analysis using clusterProfiler^47^. Visualizations were generated with dot plot and emap functions from the enrich plot package^48^. Computation of module scores was performed from the “SCT” Assay using Seurat’s function “AddModuleScore” which utilizes the method by Tirosh^49^. Pairwise inference statistics within age groups were performed using non-parametric Wilcoxon tests with p-value adjustment for multiple testing by Holm. Visualization of single-cell data was performed with build Seurat functions, module score results and differential gene expression in volcano plots was done with ggplot2^50^ and (log2FC) of candidate genes in heatmaps have been performed with the package “ComplexHeatmap”^51^.

### Gene Expression Omnibus Data Collection and Analysis

We leveraged the Gene Expression Omnibus (GEO), a comprehensive public repository that amasses high-throughput sequencing and microarray datasets from global research entities, for our investigation. A systematic search was performed utilizing the keywords "Homo sapiens", “T1D”, and “T2D” to pinpoint relevant expression datasets for our analysis. This search yielded two pertinent datasets, GSE164416 and GSE228267. Differential gene expression analysis between “healthy controls” and “patients” samples was executed using GEO2R, a user-friendly online utility that facilitates the comparison of datasets within a GEO series to identify differentially expressed genes under varying experimental paradigms. This analysis was carried out by employing the DESeq2 package, which utilized NCBI’s computed raw count matrices as input. Statistical significance was ascertained through the adjusted P-value, applying the Benjamini-Hochberg method for controlling the false discovery rate. Criteria for statistical significance were set at a fold change threshold greater than 0.5 and an adjusted P-value of less than 0.05.

### Statistical Analysis

The results were analyzed through the statistical program GraphPad Prism 9.0 (San Diego, CA, USA). The data were expressed as mean ± standard error of the mean (SEM) and D’Agostino and Pearson test was used to verify the normality of the samples, followed by the unpaired t-test for the comparison between AL and IF groups in all aged groups. Other data sets were tested for normality using the Kolmogorov-Smirnov test followed by a one-way ANOVA analysis of variance with Bonferroni multiple comparisons as post-test. The minimum significance level established was *P* < 0.05.

### Contact for reagent and resource sharing

Further information and requests for reagents and resources shall be addressed to the Lead Contact, Prof. Dr. Stephan Herzig.

### Data and Software Availability

Raw data files for the RNA sequencing and analysis will be deposited as a Series in the NCBI Gene Expression Omnibus.

## Supplementary Material

**Supplementary figure 1.**
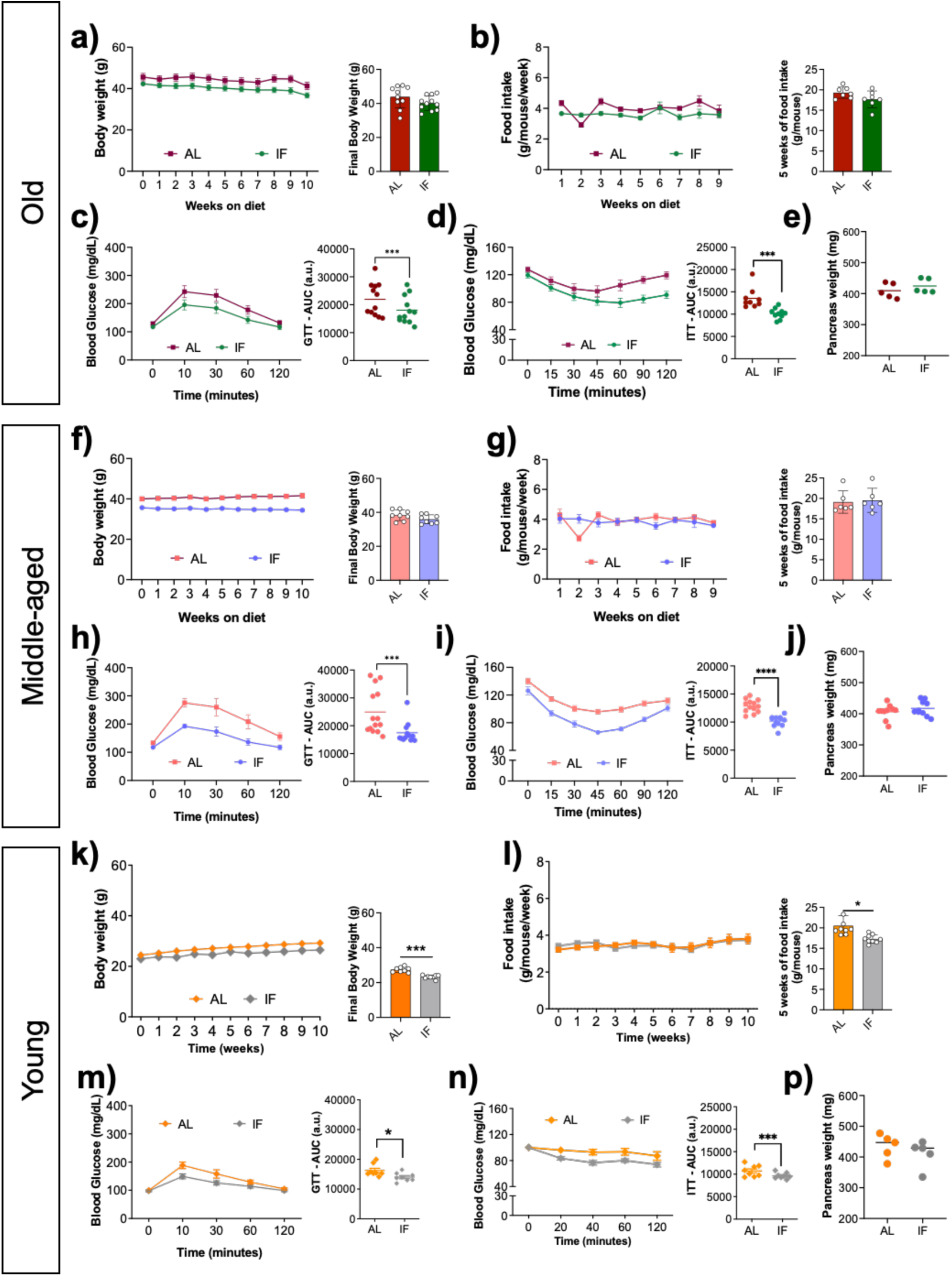
A short period of intermittent fasting intervention improves glucose homeostasis in all ages. Body weight during and at the end of 5 weeks of dietary intervention (Old: a, Middle-aged: f, and young: k); Weekly and cumulative food intake (Old: b, Middle-aged: g, and young: l); Glucose tolerance test (2.5 mg/kg) after 5 weeks of IF intervention (Old: c, Middle-aged: h, and young: m); Insulin tolerance test (0.8 U/kg) after 5 weeks of IF intervention (Old: d, Middle-aged: i, and young: n); Pancreas Weight (Old: e, Middle-aged: j, and young: o). AL: Ad libitum; IF: Intermittent fasting; GTT: Glucose tolerance test; ITT: Insulin tolerance test and AUC: Area under curve. The data were expressed as the mean ± standard error of the mean (Old: n = 9/group; middle-aged: n = 8/group; young mice: n= 8/group). Statistical differences,*p<0.05, ***p <0.001.

**Supplementary Figure 2.**
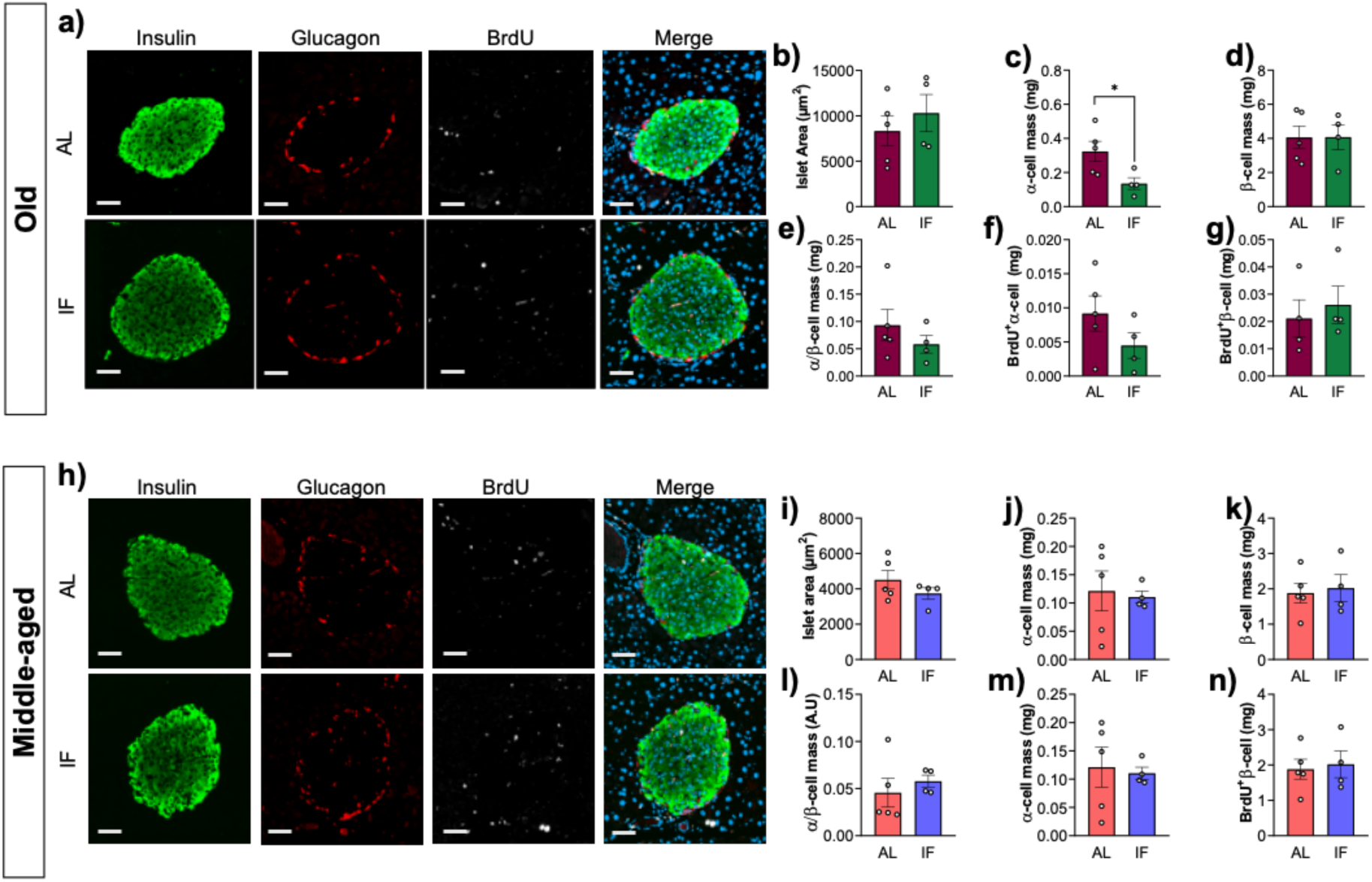
Long Intermittent fasting intervention does not affect islet morphology in old and middle-aged mice. Immunofluorescence from Isolated Islets from old- and middle-aged mice exposed to 10 weeks of intermittent fasting (a, h); Islet area μm^2^ (b, i); α-cell mass (c, j); β-cell mass (d, k); α- and β- cell ration (e, l); mass of α- and β-cells positively stained for BrdU – proliferation (old: f and g; young m and n, respectively). Insulin positive cells: Green; Glucagon positive cells: Red; BrdU positive cells: White; and Nuclear staining (DAPI): Blue – Scale bar 100μm. AL: Ad libitum; IF: Intermittent fasting. The data is expressed as the mean ± standard error of the mean (n=5 in the AL group and 3 in the IF group). Statistical differences,*p<0.05, **p <0.01.

**Supplementary Figure 3.**
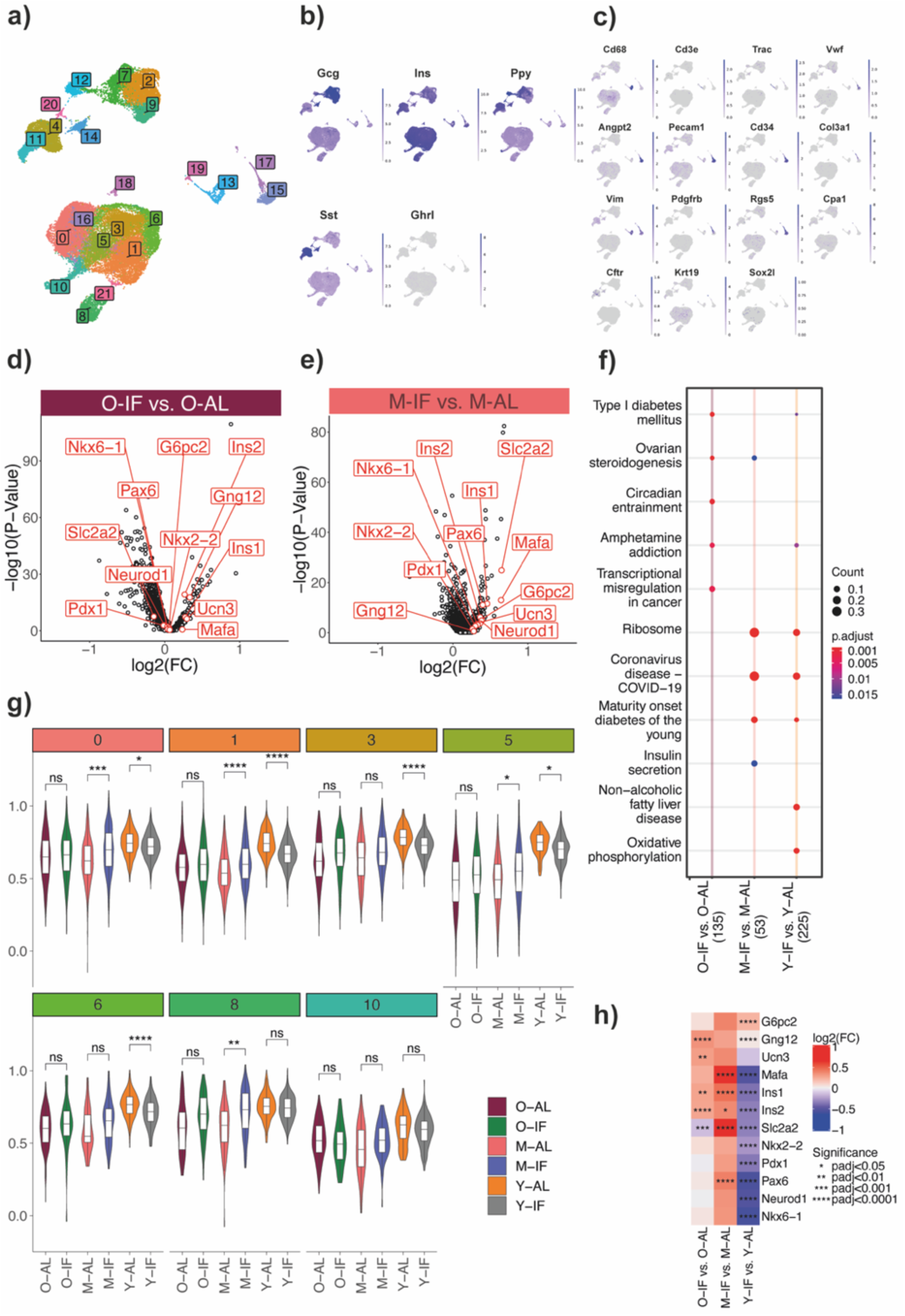
Young mice exposed to long IF intervention present reduced transcriptomic activity scores for maturation and development also in subpopulations of β-cell. UMAPs of scRNA-seq data indicating 22 statistical clusters (a), and expression of main markers for endocrine (b) as well as non-endocrine (c) cell types. Results of differential gene expression analysis upon IF vs. AL in old (d) and middle old (e) animals. Volcano plots show biological effect size (log2(FC) on the x-axis and statistical significance (-log10(P-value) on the y-axis. Comparison of over-representation analysis results of KEGG pathways in differentially expressed genes between all age groups (f). Comparison of module scores for beta cell maturation markers between IF and AL in all age groups for all beta cell clusters (g). Beta cell clusters with only a few cells have been excluded from this analysis (Clusters 16, 18 21). Heatmap of differential gene expression results (log2(FC)) of feeding invention for all age groups (h). Feeding regimen-based experimental groups are intermittent fasting (IF) vs ad libitum (AL) in old (O-IF vs. O-AL), middle-aged (M-IF vs. M-AL), and young animals (Y-IF vs. Y-AL). Statistical significance is indicated with asterisks. * padj<0.05; ** padj <0.01; *** padj<0.001; **** padj<0.0001 in (g) and (h).

